# Quantifying Crop Disease Trait Dynamics through Longitudinal Imaging and Temporal Analytics

**DOI:** 10.64898/2026.08.14.744665

**Authors:** Amanda Ewen, Rodrigo Godoy Mendez, Karar Al-Shanoon, Dawn Omoluabi, Anjana Samarasinghe, Maria Alejandra Oviedo-Ludena, Karina Chimbo Huatatoca, Kara Glor, Keiko Nabetani, Randy Kutcher, Lipu Wang, Ian Stavness, Lingling Jin

## Abstract

Reliable and objective phenotyping is essential for plant breeding programs to characterize genetic variation and accelerate crop improvement. Conventional disease assessment relies on expert visual scoring, which is labor-intensive, subjective, and prone to inter- and intra-rater variability. Although image-based phenotyping methods have been proposed, many require manual intervention, specialized imaging setups, or single time-point measurements, limiting their ability to capture disease progression over time. Here, we present a pipeline for longitudinal plant disease phenotyping that quantifies wheat stripe rust and leaf rust progression from time-series images. The pipeline performs semi-automated leaf and automated pustule segmentation from images acquired *in situ*, enabling objective disease severity estimation with minimal user intervention and without requiring solid backgrounds or manual leaf manipulation or detachment. By extracting temporal traits, including disease severity trajectories and standardized area under the disease progress curve, the method provides a comprehensive characterization of disease development throughout infection. Association between automated and expert assessments was moderate for stripe rust (*R*^2^ = 0.58) and strong for leaf rust (*R*^2^ = 0.85), while expert inter-rater reliability was moderate for both diseases (ICC = 0.675 and 0.800, respectively). The proposed approach establishes a scalable and reproducible framework for longitudinal disease phenotyping in controlled environments, with broad applications in disease resistance screening and crop breeding.

## 1. Introduction

Effective and reliable plant phenotyping can support modern genetic crop improvement. Traditionally, phenotyping in plant breeding programs relies heavily on manual visual scoring by trained experts. For example, disease severity in crops such as wheat or canola is often assessed through ordinal scales based on visible symptoms. While expert scoring provides valuable domain knowledge, it is inherently time-consuming, subjective, and difficult to standardize across observers and experimental conditions. Variability between scorers, inconsistencies across time, and limited temporal resolution reduce the reproducibility and scalability of such measurements. Furthermore, because these assessments are typically recorded as final scores without preserving the underlying raw observations (e.g., images, detailed measurements), it is often impossible to revisit or re-evaluate the data if anomalies, errors, or unexpected patterns arise. This lack of traceability and auditability further limits data reliability. Collectively, these limitations are amplified in large-scale breeding programs where many genotypes must be evaluated across multiple environments and time points.

In this paper, we present a computational pipeline for quantifying dynamic crop disease progression through longitudinal imaging and temporal analytics. Applied to wheat disease experiments for stripe rust and leaf rust, the pipeline leverages high-frequency hourly image acquisition, semi-automatic plant segmentation, and automatic disease-area segmentation to continuously track disease development and quantify disease severity throughout infection. The resulting time-series measurements enable reproducible disease progression curves and biologically meaningful temporal traits. Comparison with assessments from three independent expert raters demonstrates that the proposed approach provides accurate, objective, and reproducible estimates of disease severity while eliminating the subjectivity and limited temporal resolution associated with conventional visual scoring methods.

Overall, the proposed pipeline establishes a scalable computational approach for time-resolved plant phenotyping, integrating automated imaging, plant organ segmentation, and quantitative disease analysis into a unified workflow. Beyond wheat rust monitoring, our method provides a general approach for modeling dynamic plant traits from time-series images, with applications in disease phenotyping, stress response analysis, and data-driven crop improvement.

## 2. Related Work

Wheat rust diseases are among the most economically significant fungal infections affecting wheat, causing substantial yield and quality losses. Disease severity is typically quantified as the percentage of infected tissue, estimated as the proportion of leaf area covered by pustules or lesions (1). Monitoring severity over time enables characterization of disease progression using metrics such as the Area Under the Disease Progress Curve (AUDPC) (2) and the Area Under the Disease Progress Stairs (AUDPS) (3).

Traditional assessment relies on manual estimation of diseased leaf area using standard rating scales (1). This process is labor-intensive, time-consuming, and subjective, leading to variability among evaluators. Automated approaches often classify disease severity into discrete categories defined by these scales, but the number of classes varies widely across studies, from 3 to 10 levels, depending on the assessment protocol (4–16). While useful for distinguishing stages, these labels provide only discrete snapshots and do not capture continuous disease progression, even when applied over time.

Previously proposed methods for automated severity estimation include: feature extraction with classification (17– 20), thresholding-based approaches (21), color-based segmentation (22), classical machine learning (23), clustering and quantum-based methods (24), and deep learning(19, 20, 25, 26).

Despite these advances, several limitations remain. Some methods analyze only small leaf regions (21, 24), although disease distribution is often heterogeneous. Whole-leaf approaches frequently require manual handling, standardized backgrounds, or controlled imaging setups (17–20, 22, 23, 25), limiting scalability. Many also require leaf detachment (21, 22), preventing repeated measurements and longitudinal analysis. As a result, most approaches focus on single timepoint assessment and do not support temporal metrics. Recent longitudinal approaches (19, 20) address some of these issues but rely on structured acquisition protocols with standardized backgrounds, markers, and require repeated human intervention, reducing applicability in natural or large-scale settings.

## 3. Materials and Methods

In this study, we develop a computational pipeline to quantify the temporal progression of wheat foliar diseases from longitudinal image sequences. The unit of analysis is an individual wheat flag leaf captured repeatedly through high-frequency time-lapse imaging, enabling passive longitudinal monitoring with only a fixed imaging device and minimal user intervention once configured. The pipeline employs computer vision and image processing techniques to analyze the temporal image sequences, enabling consistent and objective measurement of disease development over time. It also generates visualizations of colour changes over time, revealing temporal patterns of disease progression and plant stress at the individual leaf level, supporting downstream breeding program decisions and facilitating the evaluation of disease response trajectories.

The modular analysis pipeline, as shown in Figure 1, comprises leaf segmentation, disease identification, quantitative assessment of infection, and phenotype visualization.

**Figure 1.**
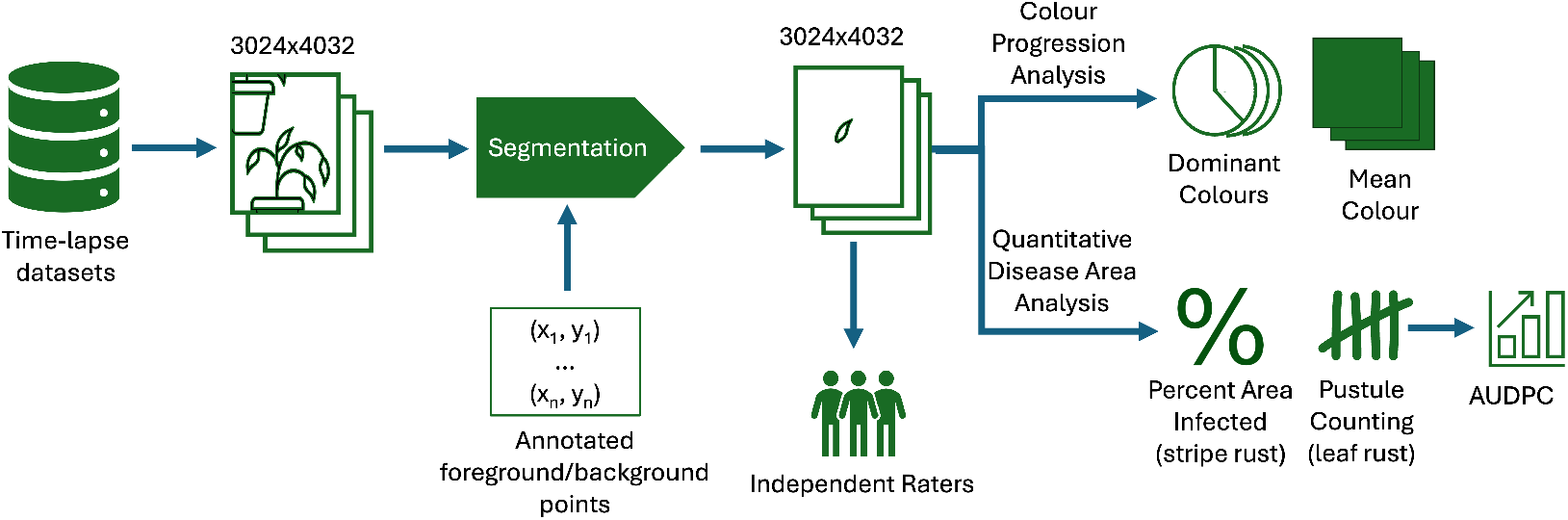
The overview of the analysis pipeline. Starting with time-series images, each image undergoes semantic segmentation followed by automatic qualitative and quantitative analysis methods. A subset of segmented images are also provided to independent expert raters for manual disease rating to validate the automatic disease ratings.

### 3.1 Experimental Set Up

Four wheat varieties with differing levels of stripe rust resistance were grown under speed breeding conditions: Avocet (susceptible), Carberry (moderately resistant), Lillian (resistant), and Avocet+Yr15 (resistant). Each variety was grown in three biological replicates, with two pots per replicate and two plants per pot. Plants received standard fertilization throughout the experiment, and growth stages were monitored using the GreenSkEye platform (27–30), together with manual observations.

Stripe rust was induced by spray inoculating flag leaves at the swollen boot to early heading stages (Z45-Z51). Following inoculation, plants were transferred to a misting chamber and maintained under environmental conditions favorable for stripe and leaf rust development (22 h light/2 h dark; 17°C/10°C day/night). After disease evaluation, standard speed breeding conditions (22h light/2h dark; 21°C/17°C day/night) were restored.

Images were captured of just two varieties (Avocet and Avocet+Yr15) daily in an initial stripe rust pilot experiment, and then expanded to capture all four varieties hourly in the subsequent experiment. Little change was observed within one day, so for consistency, only one image per day was used for analysis. An example image is shown in Figure 2. The datasets from these stripe rust experiments were combined for downstream analysis.

**Figure 2.**
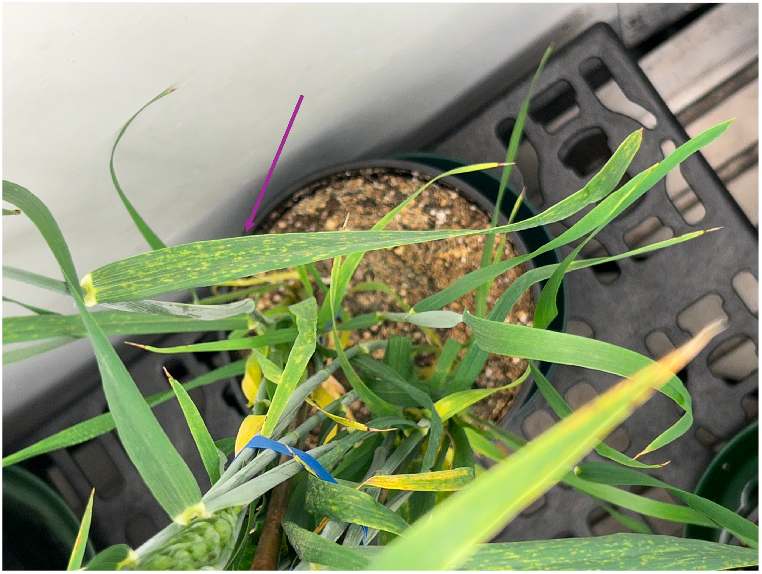
Example image of an Avocet wheat plant captured in a growth chamber using the GreenSkEye iPhone application. The arrow, added for this figure, points to the flag leaf of interest.

A further study was initialized with multiple replicates of Avocet and Avocet+Yr15 being imaged, grown and inoculated as described above. Incidentally, these plants developed leaf rust symptoms, allowing the analysis pipeline to be adapted to another disease. With unknown inoculum composition for leaf rust, the pipeline is instead tested against manual ratings.

### 3.2 Flag Leaf Semantic Segmentation

The captured images contain complex backgrounds, including surrounding vegetation, pots, and imaging equipment. In addition, the target flag leaf is occasionally partially occluded, causing it to appear as multiple disconnected components. To robustly isolate individual flag leaves under these conditions, we employ an adapted version of the Segment Anything Model (SAM) (31), which generates segmentation masks from user-provided prompt points.

Segmentation prompts were automatically propagated across longitudinal image sequences, eliminating the need for manual point selection in each image. Because consecutive images typically exhibit only minor changes in leaf position and appearance, the propagated prompts generally remained valid. Manual reselection was required only when substantial leaf movement or occlusion prevented accurate segmentation. All samples were checked for segmentation failures and none were found.

### 3.3 Colour Progression Analysis

Once images are segmented to isolate flag leaves, we made qualitative analyses using two colour-based techniques. Both techniques allow for visualization of how the leaves change over time.

#### 3.3.1 Mean Colour

For each time point *t*, the mean colour of the segmented flag leaf is computed by averaging the pixel intensities of the red (*R*), green (*G*), and blue (*B*) channels. For example,

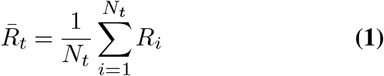

where *N*_*t*_ is the number of pixels in the segmented flag leaf at time *t*, and *R*_*i*_ is the red-channel intensity of the *i*-th pixel. Then, the mean values of the three colour channels, 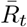, 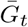, and 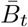, are combined to form an RGB colour vector, representing the mean colour of the flag leaf at time *t*.

#### 3.3.2 Dominant Colours

We apply the K-means clustering algorithm to the RGB pixel values of the segmented flag leaf image, partitioning the pixels into two clusters using the k-means++ initialization method (32). The resulting cluster centroids represent the two dominant colours present in the leaf at time *t*, while the proportion of pixels assigned to each cluster quantifies their relative abundance. If a leaf is diseased with rust, k=2 allows separation of healthy tissue and rust pustule colours. The temporal changes in the hue and proportion of these dominant colours provide a simple yet informative representation of disease progression and leaf senescence for qualitative analysis.

### 3.4 Quantitative Disease Area Analysis

To quantitatively assess rust severity, we measured the proportion of the flag leaf covered by pustules for stripe rust and the number of individual pustules detected per image for leaf rust. These metrics were computed independently at each time point, enabling longitudinal analysis of disease progression.

#### 3.4.1 Pustule Coverage

Input images are represented in the RGB colour space and normalized to the range [0, 1]. We first compute the *key* (black) component *K* from the CMYK colour space for each pixel:

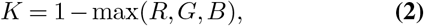

where *R, G*, and *B* denote the red, green, and blue intensity values, respectively.

Using the key component and the green channel, we then derive the magenta component (*M*), which enhances contrast between healthy tissue and rust pustules:

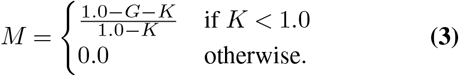

To separate diseased from healthy tissue, we apply Otsu’s thresholding method. Rust pustules exhibit elevated magenta values, therefore pixels with *M* > *T*, where *T* is the computed threshold, are classified as diseased.

Finally, disease severity for each image is quantified as the percentage of leaf area occupied by the detected pustules, referred to as pustule coverage (*P*_*t*_) at time *t*:

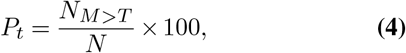

where *N*_*M*>*T*_ is the number of pixels exceeding the threshold and *N* is the total number of pixels within the segmented flag leaf at time *t*.

#### 3.4.2 Pustule Counting

For leaves infected with leaf rust, we further characterize disease progression by quantifying individual pustules. Starting from the thresholded binary mask, we apply morphological closing followed by opening operations to smooth the segmentation and remove small artifacts. Connected components larger than (*k*) pixels (*k* = 5, 000, determined empirically for this dataset) are then excluded, as they are unlikely to correspond to individual pustules. Finally, the remaining connected components are counted to estimate the total number of leaf rust pustules present on the leaf surface.

### 3.5 Manual Ratings

Manual assessments of stripe rust and leaf rust severity were performed independently by three expert raters using a modified Cobb scale (1), which is capped at 37% because the maximum proportion of internal leaf tissue occupied by erupting pustules is biologically limited to approximately 37% (1). Segmented images of 16 stripe rust leaves and 18 leaf rust leaves at two time points, were scored and used as reference values for comparison with the automated disease estimates. To assess intra-rater reliability, five images from each disease type were randomly selected and repeated three times within the evaluation dataset. Consequently, each rater provided 42 severity ratings for stripe rust and 46 for leaf rust. Intra-rater and inter-rater reliability were quantified using two-way random-effects (inter-rater) or two-way mixed-effects (intra-rater) intraclass correlation coefficient (ICC) models based on single-measures and the absolute agreement definition (33).

## 4. Results

### 4.1 Colour Progression Analysis

Results from mean and dominant colour analyses for representative stripe rust-infected, leaf-rust infected, and healthy leaves are presented in Fig. 3. These examples illustrate distinct temporal colour progression patterns associated with disease development and show differences between infected and healthy leaves over time.

**Figure 3.**
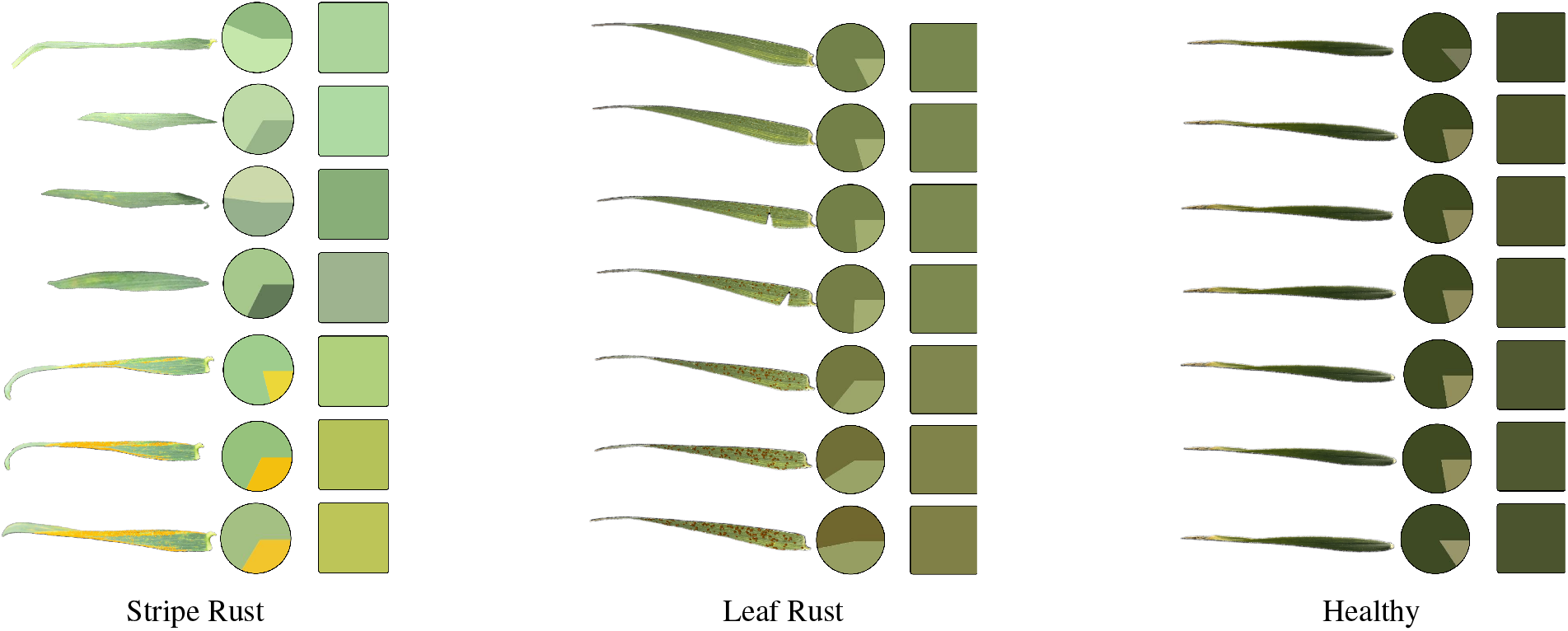
Example Colour Progression analysis over two weeks for different disease conditions. The segmented leaf and its corresponding dominant colours and mean colour are shown for every two days. Variations in brightness are due to background illumination.

### 4.2 Quantitative Disease Progression

#### 4.2.1 Stripe Rust

Comparing the automated disease severity estimates based on pustule coverage with the mean expert ratings yielded a Mean Squared Error (MSE) of 103.76, a Root Mean Squared Error (RMSE) of 10.19, and an *R*^2^ value of 0.58 (Fig. 4, left). The only leaf without visible stripe rust pustules was assigned an estimated severity of 6.7%. Fig. S1 compares automated estimates with ratings from individual experts.

**Figure 4.**
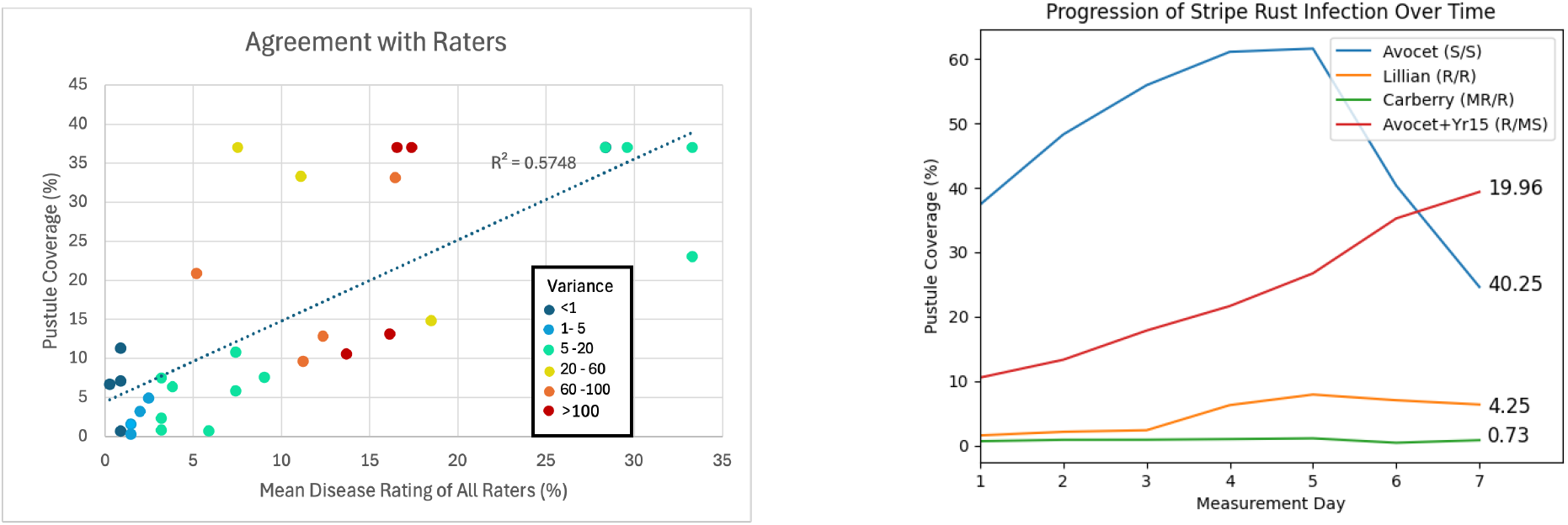
Left: Automated disease severity estimates derived from pustule coverage versus mean expert ratings for each stripe rust sample. Points are coloured according to sample variance between three expert ratings. Right: Temporal disease progression in representative wheat leaves for resistant, moderately susceptible, and susceptible leaves are plotted with AUDPC values on the right-hand side. The legend denotes variety name and (expected/expert-rated) disease severity (S=susceptible, M=moderately, R=resistant).

Temporal disease progression, including pustule coverage and standardized AUDPC (normalized by the number of observations), are presented for representative resistant, moderately susceptible and susceptible leaves in Fig. 4 (right).

#### 4.2.2 Leaf Rust

Comparing the automated disease severity estimates based on pustule count with the mean expert ratings yields an *R*^2^ value of 0.853, as shown in Fig. 5 (left). Healthy leaves showing no leaf rust symptoms had an average of 3±2 pustules when automatically counted. Fig. S2 compares automated estimates with ratings from individual experts.

**Figure 5.**
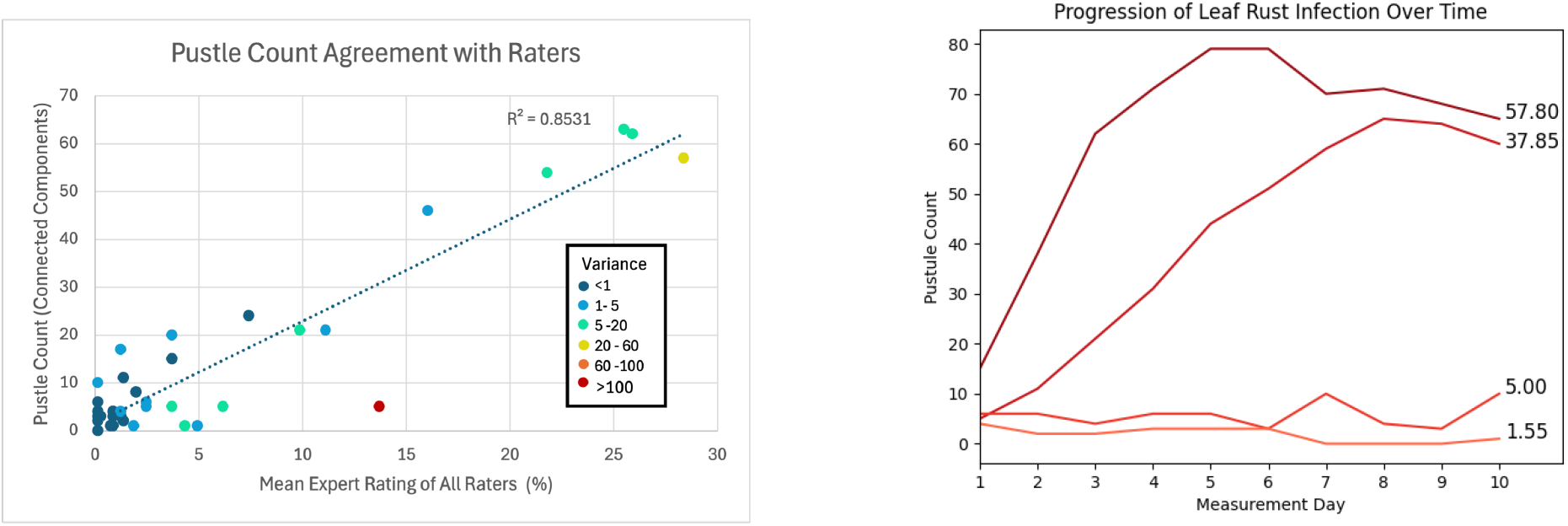
Left: Automated disease severity estimates derived from pustule count versus mean expert ratings for each leaf rust sample. Points are coloured according to sample variance between three expert ratings. Right: Temporal progression of pustule counts in four representative flag leaves. The red color gradient reflects expert disease severity ratings, ranging from light red (low severity) to dark red (high severity). The AUDPC is shown to the right of each curve.

Temporal progression of pustule counts for representative leaves identified as resistant or susceptible and their corresponding AUDPC values are shown in Fig. 5 (right).

### 4.3 Intra- and Inter-Rater Reliability

Intra-rater reliability across diseases and raters are shown in Table 1. However, the relatively wide 95% confidence intervals indicate uncertainty in these estimates.

**Table 1.**
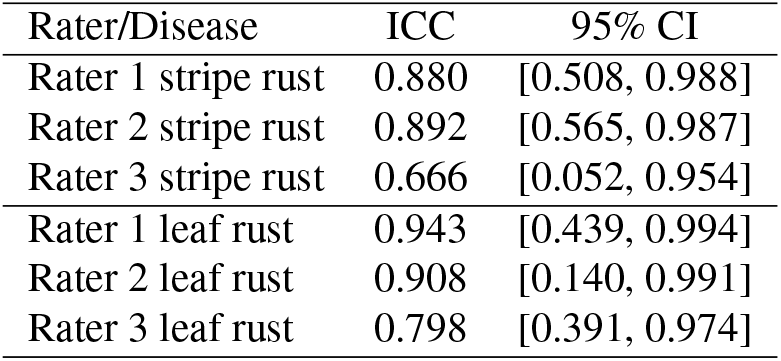
Intra-rater reliability for disease severity assessments measured by the intraclass correlation coefficient (ICC) and corresponding 95% confidence intervals for stripe rust and leaf rust.

| Rater/Disease | ICC | 95% CI |
| --- | --- | --- |
| Rater 1 stripe rust | 0.880 | [0.508, 0.988] |
| Rater 2 stripe rust | 0.892 | [0.565, 0.987] |
| Rater 3 stripe rust | 0.666 | [0.052, 0.954] |
| Rater 1 leaf rust | 0.943 | [0.439, 0.994] |
| Rater 2 leaf rust | 0.908 | [0.140, 0.991] |
| Rater 3 leaf rust | 0.798 | [0.391, 0.974] |

Inter-rater reliability ICC scores for both diseases, computed across all three raters, are shown in Table 2. In addition, the variance of the ratings for each sample was calculated to quantify the level of disagreement between raters (Figures 4-left and 5 - left), and the agreement between the automated method and individual expert raters is shown in Fig. S3.

**Table 2.** Inter-rater reliability for disease severity assessments measured by the intraclass correlation coefficient (ICC) and corresponding 95% confidence intervals for stripe rust and leaf rust.

| Disease | ICC | 95% CI |
| --- | --- | --- |
| Stripe rust | 0.674 | [0.376, 0.833] |
| Leaf rust | 0.800 | [0.682, 0.854] |

## 5. Discussion

Temporal changes in mean and dominant leaf colour provide an intuitive visualization of disease progression. For stripe rust, disease development is characterized by the increasing appearance of yellow hues in the dominant colour profile and a corresponding shift in mean colour away from healthy green. In contrast, early-stage leaf rust is less distinguishable by colour, but increasing disease severity produces a gradual transition toward desaturated brown tones associated with expanding pustules. Healthy leaves maintain relatively stable colour profiles over the observation period. Cameras are set to fixed white balance within a growth chamber environment with fixed lighting, so colour calibration was not used. In scenes with lighting variation colour calibration may be used.

Intraclass correlation coefficient analysis highlights the subjectivity of expert disease severity assessments from longitudinal image data. Intra-rater reliability ranged from 0.798 to 0.943 for leaf rust and from 0.666 to 0.892 for stripe rust, indicating greater variability in repeated assessments of stripe rust. The associated 95% confidence intervals were wide for several raters, particularly for stripe rust, suggesting substantial uncertainty in reliability estimates and further emphasizing the challenges of consistent visual disease assessment. Inter-rater reliability was also poor to moderate, with ICC values of 0.682 to 0.854 for leaf rust and 0.376 to 0.833 for stripe rust, demonstrating that different experts may assign substantially different scores to the same sample. These results underscore the limitations of traditional visual rating methods, particularly for stripe rust, where disease patterns can be difficult to interpret consistently. The observed variability motivates the use of automated phenotyping approaches that provide more objective, consistent, and reproducible estimates of disease severity.

Comparison of the automated pustule coverage estimates with the mean expert ratings showed the strongest agreement at low and high disease severity, with greater discrepancies at intermediate levels. Overall, the severity estimates from the automated method showed moderate association with experts as indicated by an *R*^2^ of 0.58. Variability within expert ratings may partly explain moderate amount of association. Because measurements were collected from the same leaves at two time points, some within-leaf correlation may be present, and the reported *R*^2^ may be slightly inflated. Our use of *R*^2^ follows from previous work (25). Pairwise ICC values also indicate variability between raters and moderate agreement with automatic stripe rust disease estimates (Fig. S3).

Many of the largest discrepancies between automated and manual ratings were associated with the presence of telia, a late-stage rust structure that appears as flat brown lesions rather than bright pustules. For example, the leaf in Fig. 6 received expert ratings ranging from 1 (0.37%) to 70 (25.9%), whereas the automated method estimated the maximum severity of 37%. Because telia exhibit low green intensity, they are often classified as diseased tissue by the automated method. In contrast, expert raters differed in whether telia should be included in disease severity estimates, leading to substantial disagreement. These results highlight the need for standardized, objective approaches to disease severity quantification.

**Figure 6.**
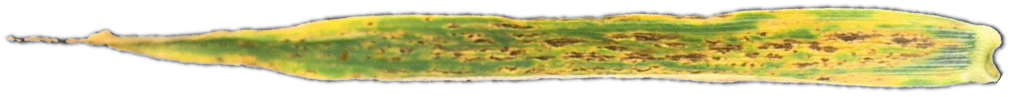
Segmented flag leaf infected with stripe rust where rust disease appears as telia rather than traditional pustules.

The automated pustule counting method for leaf rust showed strong agreement with expert ratings, achieving an *R*^2^ of 0.85. Inter- and intra-rater reliability were also higher for leaf rust than for stripe rust, indicating greater consistency among expert assessments. This strong correspondence suggests that automated pustule counting provides an objective and reliable measure of leaf rust severity.

One sample exhibited unusually high rating variance (Fig. 7) because it displayed symptoms of both leaf rust and stripe rust. Consequently, one expert assigned the maximum severity rating (37%), whereas another rated it at only 0.37%. The two automated methods captured different aspects of this mixed infection: pustule counting detected only five discrete pustules and produced a low severity estimate, while pustule coverage assigned the maximum severity.

**Figure 7.**
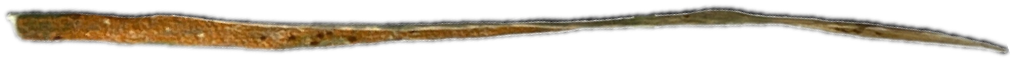
Segmented flag leaf showing symptoms of both leaf and stripe rust.

This example demonstrates that the two automated approaches capture complementary disease characteristics. Pustule counting is sensitive to discrete lesions, whereas pustule coverage reflects more diffuse or extensive symptoms. Together, these methods provide a more comprehensive and objective characterization of disease severity than manual assessment alone.

AUDPC captures disease development over time and clearly distinguishes resistant, moderately resistant, and susceptible samples for both stripe rust and leaf rust, regardless of whether disease severity is quantified by pustule coverage or pustule count. Longitudinal AUDPC analysis also captures temporal dynamics that single time-point measurements may miss. For example, in Fig. 4 (right), disease severity peaks before the final observation as infected leaves begin to develop chlorosis. Relying only on the final assessment could therefore underestimate disease severity, whereas AUDPC provides a more comprehensive measure by integrating disease progression over time.

The proposed method has only been tested in controlled growth chamber environments, where stable lighting conditions support reliable colour analysis and disease detection. Leaf curling and movement can also affect segmentation and disease quantification by altering leaf orientation. This variability is partially mitigated by frequent image acquisition, which enables robust temporal analysis and facilitates visual inspection of longitudinal image sequences. A remaining bottleneck is the segmentation step, which currently requires user-selected foreground and background points to isolate flag leaves. This manual interaction limits scalability, making full automation of the segmentation process an important direction for future work.

## 6. Conclusion

This work presents a computational method for automated, longitudinal quantification of wheat stripe rust and leaf rust disease progression from time-series imagery. By integrating semi-automated leaf and automated pustule segmentation, and temporal analytics, the method enables objective measurement of disease severity and progression with limited manual intervention, while preserving the complete image record for future reanalysis. Validation against expert assessments demonstrated agreement with manual ratings while highlighting the variability among human raters, underscoring the value of standardized and reproducible image-based phenotyping.

Beyond single time-point severity estimates, our method captures rich temporal phenotypes, including disease progression curves and standardized AUDPC, providing a more comprehensive characterization of host-pathogen interactions throughout disease development. To the best of our knowledge, this is the first semi-automated pipeline for wheat stripe rust and leaf rust phenotyping that quantifies disease severity directly from images acquired *in situ*, without requiring solid backgrounds, manual leaf manipulation, or other specialized imaging preparations. By enabling high-throughput and reproducible disease assessment under realistic imaging conditions, this work provides a practical foundation for digital plant pathology screening and has potential applications in crop breeding, disease resistance evaluation, and precision agriculture.

## Code and Data Availability

Software and datasets are available at:

https://github.com/USask-BINFO/greenskeye_analysis and https://greenskeye.usask.ca/speedbreeding/.

### Acknowledgments

This work was funded by the Natural Sciences and Engineering Research Council of Canada, Canada Foundation for Innovation, and Saskatchewan Ministry of Agriculture.

## Conflict of Interest

The authors declare no conflict of interest.

## Author Contributions

A.E., L.J., I.S., L.W., and R.K. conceived and designed the study. A.E. developed the analytical pipeline. A.E., R.M., K.A., D.O., and L.J. collected the imaging data, R.M., D.O., and A.S. organized and managed the data, while M.A.O., K.C.H., K.G., L.W., and R.K. conducted the plant experiments. A.E. prepared the manuscript. L.J., and I.S. supervised the research and revised the manuscript.

## Supplementary Material

**Figure S1.**
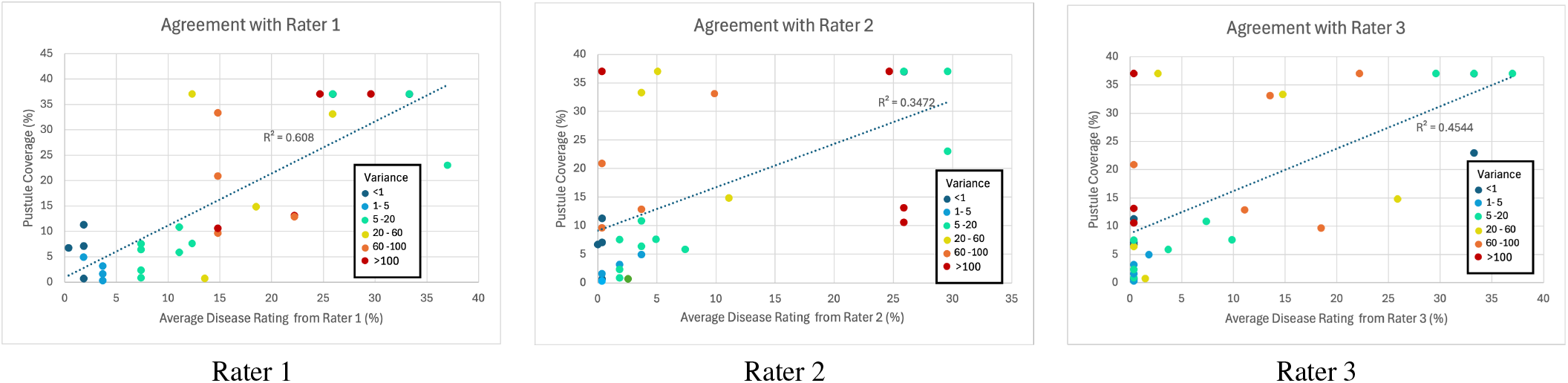
Automated disease severity estimates derived from pustule coverage vs individual expert ratings for each stripe rust sample. Points are coloured according to sample variance between three expert ratings.

**Figure S2.**
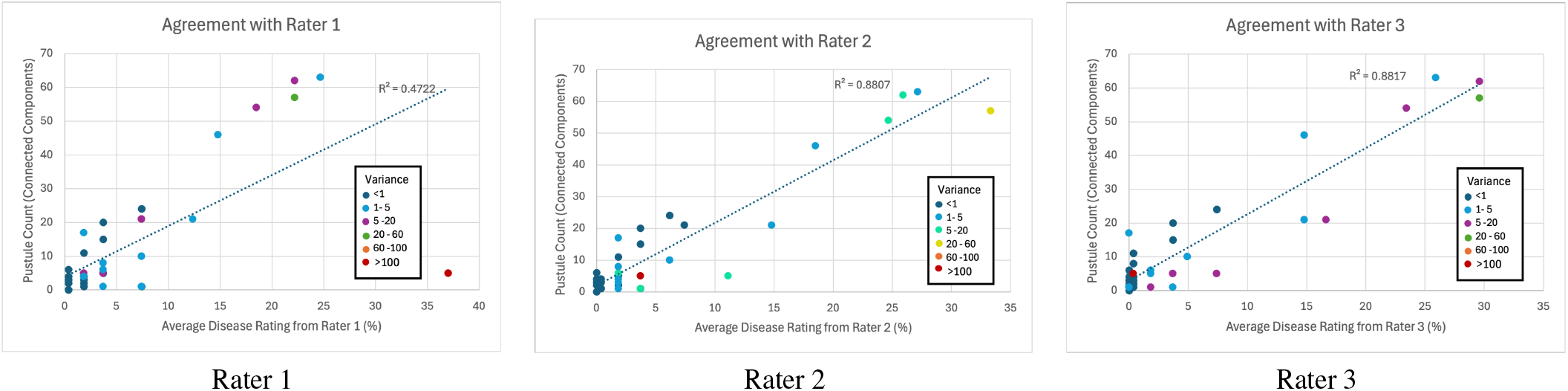
Automated disease severity estimates derived from pustule count versus individual expert ratings for each leaf rust sample. Points are coloured according to sample variance between three expert ratings.

**Figure S3.**
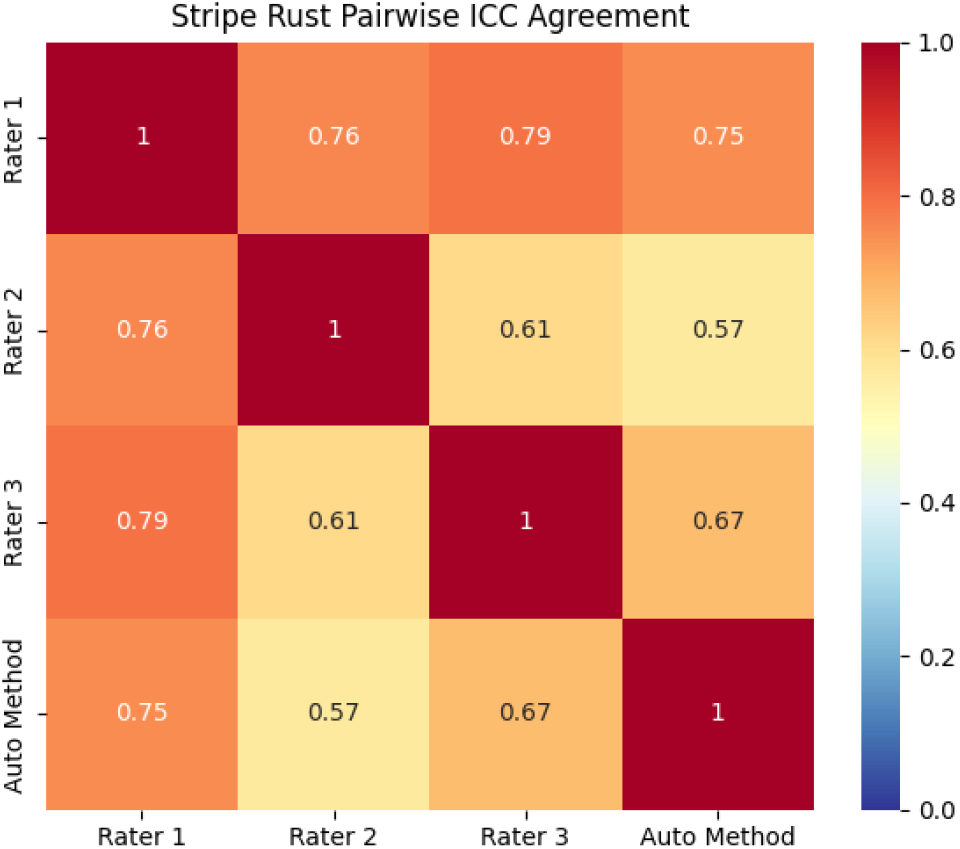
Pair-wise intraclass correlation coefficient (ICC) values between each rater and the automated pipeline for Stripe Rust severity.

## Notes

https://github.com/USask-BINFO/greenskeye_analysis

https://greenskeye.usask.ca/speedbreeding

### Competing Interest Statement

The authors have declared no competing interest.

